# Evaluation of the Safety of Percutaneous Electrical Stimulation of the Phrenic and Femoral Nerves in a Chronic Porcine Model: A GLP Study

**DOI:** 10.1101/2020.08.17.254847

**Authors:** John O’Mahony, Carlus Dingfelder, Igor Polyakov, Trace Jocewicz, Jennifer Mischke

## Abstract

**Purpose:** Diaphragm pacing has been proposed as a method to prevent ventilator-induced diaphragm dysfunction (VIDD) during mechanical ventilation (MV). The present study assessed the safety of lead deployment and control of diaphragm inspiratory work in synchrony with MV utilizing percutaneous electrical phrenic nerve stimulation (PEPNS) in a sedated and ventilated porcine model.

**Methods:** The ability to safely place PEPNS four-electrode leads near the femoral nerve using ultrasound visualization and electrical stimulation to guide lead placement using a through the needle (TTN) approach was assessed for 4 animals. The feasibility of using the PEPNS system to activate the diaphragm in synchrony with inspiration within a desired target Work of Breathing (WOB) between 0.3 and 0.7 joules/L over eight hours was tested using three of the four animals with the fourth used as a control. The ability to control WOB during inspiration was assessed for flow and pressure-controlled breaths using a flow and pressure sensor attached to the wye, the connector joining the inspiratory and expiratory limbs to the endotracheal tube. Overall health (moribundity) was assessed at baseline and throughout the study until Day 30 for the surviving animals. Gross pathology and histopathological studies were performed on the femoral nerves and diaphragm muscle tissue following termination of the animals at Day 30.

**Results:** The lower bound estimate of the proportion of successful stimulation within the desired level of WOB was 95.1%, achieving the study endpoint. Triggering synchrony was statistically significant at <88ms (p<0.0001) with WOB able to be maintained between 0.3 and 0.7 joules/L. There was no evidence of tissue or nerve damage nor impact on overall animal health associated with lead placement or electrical stimulation.

**Conclusion:** The PEPNS leads were found to be safe for their intended use and the PEPNS system met preestablished study endpoints for synchrony and stimulation efficacy.

## INTRODUCTION

Diaphragmatic weakness and atrophy have been reported to develop rapidly in patients following the initiation of mechanical ventilation (MV) with significant correlation to duration of ventilator support [1]. Even brief periods of MV may result in diaphragm weakness which can lead to Ventilator-Induced Diaphragm Dysfunction (VIDD). The development of VIDD can lead to weaning failure, contribute to prolonging weaning times, and impact patient mortality [2, 3].

Stimulation of the phrenic nerves to induce diaphragmatic movement was first reported by Sarnoff in 1948 [4]. The clinical use of implantable diaphragmatic pacemakers has subsequently been validated in patients requiring chronic, retracted MV [5]. Intermittent electrical stimulation of the phrenic nerves to pace the diaphragm has been suggested as a strategy to minimize the reduction in diaphragm atrophy and to potentially reduce weaning failure [2]. Recently, O’Rourke et al demonstrated the ability to safely and successfully place percutaneous multipolar leads in the anatomical region of the neck close to phrenic nerves in patients on mechanical ventilation and to use these leads to deliver percutaneous electrical phrenic nerve stimulation (PEPNS) [6]. The authors reported that they were able to demonstrate the feasibility of using this approach to synchronize electrical stimulation with inspiration while maintaining Work of Breathing (WOB) within defined limits.

The above clinical study was preceded by preclinical testing in a sedated and ventilated porcine model to investigate the safety of PEPNS lead deployment using ultrasound imaging, control of the diaphragm inspiratory work in synchrony with MV utilizing percutaneous electrical phrenic nerve stimulation (PEPNS) and prevention of damage to the diaphragm muscle resulting from fatigue or electrical stimulation dyssynchrony. The present preclinical study was conducted as a result of previous studies reporting acute diaphragm injury and weakness when the diaphragm contracts against an excessive load or when electrical stimulation dyssynchrony occurs and causes the diaphragm muscle to lengthen during exhalation [7–10]. Herein we report these preclinical studies which were conducted in compliance with the Food and Drug Administration Good Laboratory Practice Regulations (FDA GLP), 21 CFR Part 58; Final Rule effective September 4, 1987 and as updated where specified.

## MATERIALS AND METHODS

The present study was performed between November 2017 and February 2018 using randomly selected Yorkshire crossbreed female swine with a mean weight of 29.3 ± 3.8 kg (Table 1). All studies were carried out in an Association for Assessment and Accreditation of Laboratory Animal Care (AAALAC) approved facility and were reviewed and approved by the Institutional Animal Care and Use Committee (IACUC). The animal care protocol followed for the study is shown in S1 Table.

**Table 1.** Animal Pre-Treatment and Pre-Termination Data.

| Animal ID | Group | Gender | Treatment Date | Termination Date | Pre-Treatment Weight (kg)* | Pre-Termination Weight (kg)* | % Weight Change |
| --- | --- | --- | --- | --- | --- | --- | --- |
| 17P0923 | 01 | Female | 11/28/17 | 12/28/17 | 34.4 | 43.8 | 27.3 |
| 17P0933 | 02 | Female | 11/27/17 | 11/30/17** | 30.4 | NA | NA |
| 17P0981 | 01 | Female | 12/5/17 | 1/4/18 | 27.4 | 39.6 | 44.5 |
| 17P0986 | 02 | Female | 12/21/17 | 1/19/18 | 31.4 | 41.8 | 33.1 |
| 17P0989 | 02 | Female | 12/14/17 | 12/20/17<sup>†</sup> | 29.0 | NA | NA |
| 17P1001 | 01 | Female | 12/7/17 | 1/5/18 | 23.2 | 34.6 | 49.1 |
|  |  |  |  | Mean | 29.3 | 39.95 | 38.5 |
|  |  |  |  | SD | 3.81 | 3.96 | NA |

\* Implant and termination weights were obtained within 3 days prior to procedure date unless otherwise noted.
\*\* Early death, no pre-term weight obtained.
† Early termination, no pre-term weight obtained.
SD = Standard Deviation
Group 01 = Test; Group 02 = Control

The electrical lead used for the study was made of 4 thin, insulated wires bundled within polyurethane insulation with 4 platinum-iridium cylindrical electrodes at the distal end, spaced 8 mm apart (Fig 1). The leads were 31.5 cm in length with a diameter of 0.87 mm. Each electrode was 4.0 mm in length spaced 8.0 mm apart. Electrical stimulation was delivered by using a combination of two electrodes on the lead.

**Fig 1.**
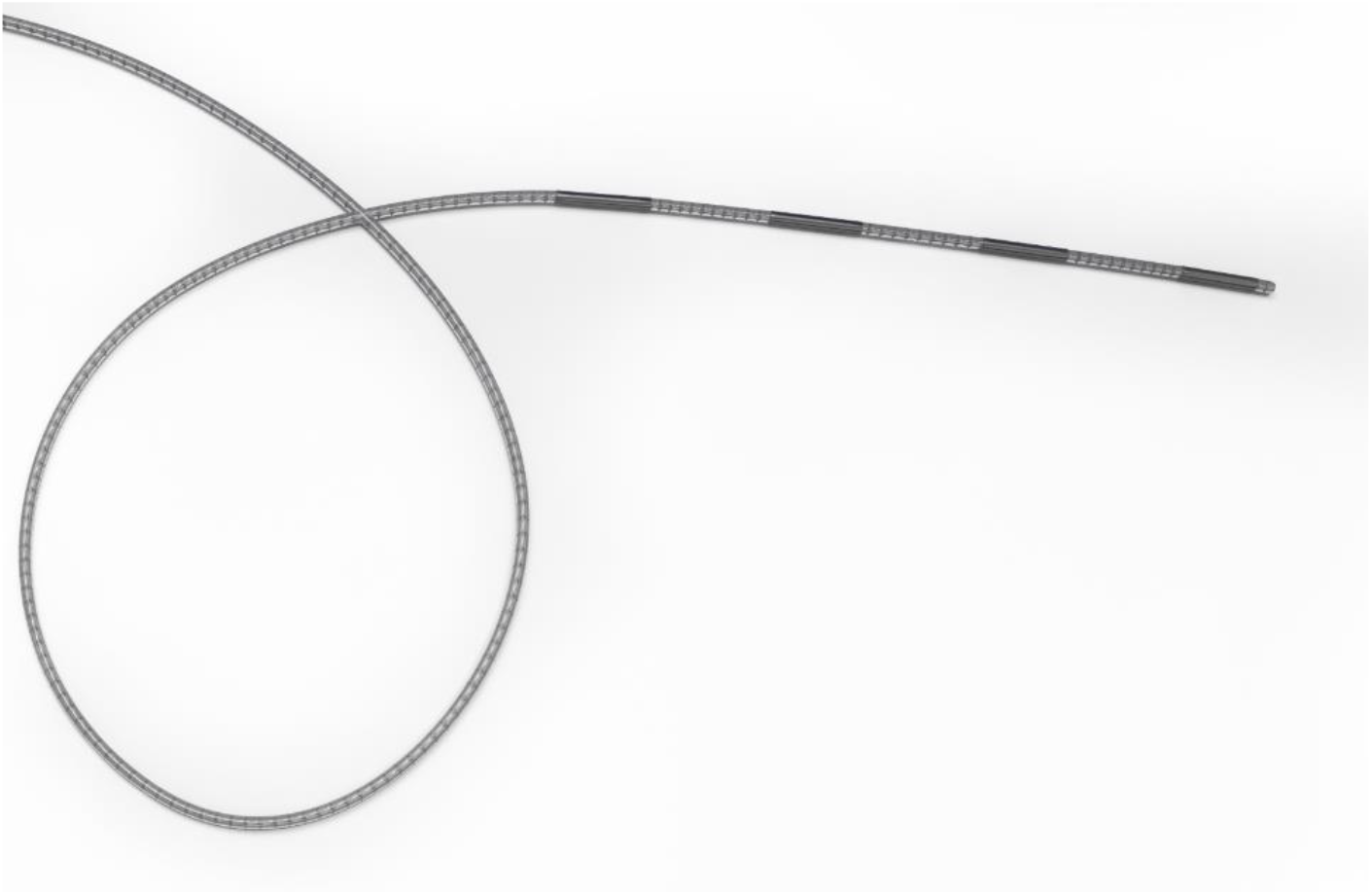
The PEPNS four electrode lead.

### Study Design

Animals were assigned in a non-randomized fashion to either a test or control arm for the study (Figs 2A and 2B). All animals had a four-pole electrode lead placed bilaterally in the femoral region near the left and right femoral nerves using ultrasound imaging to place the leads adjacent to the nerve. The test arm also had two 28G Ambu^®^ Neuroline Monopolar needles (Models 74325-36/40 and 74338-45/40, Ambu, Inc., Columbia, MD) placed bilaterally in the neck region by the thoracic inlet, near the left and right phrenic nerves in order to achieve stimulation of the diaphragm. Animals were excluded if stimulation could not be achieved in at least one leg and both phrenic nerves or if any disease or injury unrelated to the device or any treatment could have affected the study outcome. A total of six animals were utilized for this study. There were two early deaths not related to the test procedures for animals originally enrolled in the control arm.

**Fig 2.**
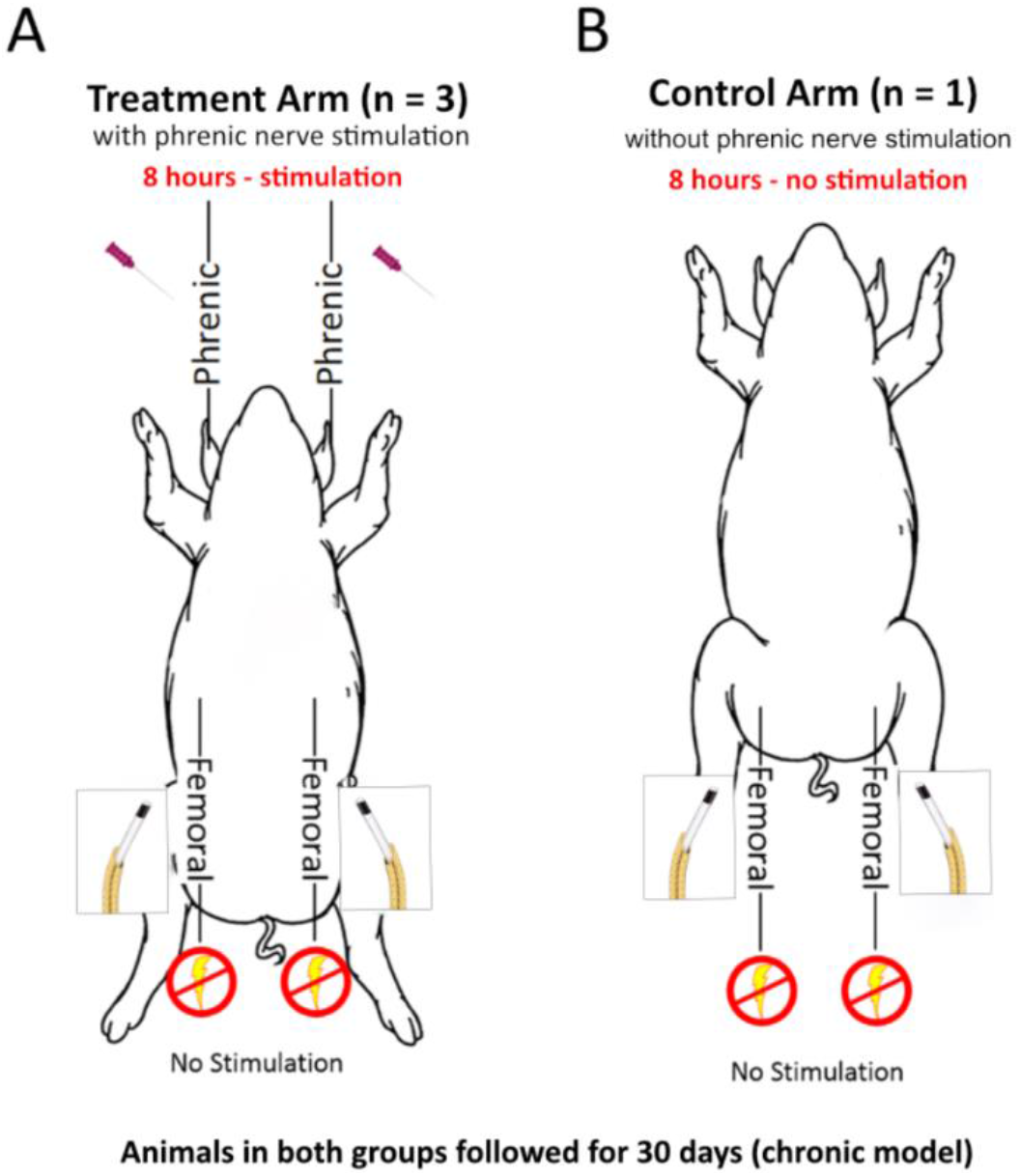
Schematic Representation of Study Design. A) treatment arm with 3 animals and B) control arm with 1 animal. Two Ambu needles were used for the bilateral stimulation of the phrenic nerves in the treatment arm. Pajunk needles were used to deploy the PEPNS Leads. Electrical symbol denotes stimulation was not continuous. Femoral nerve stimulation was tested after initial lead deployment and at 8 hours.

The primary study objectives were to confirm the safety of lead placement near the femoral nerve and the ability to safely capture the phrenic nerve with associated electrical stimulation of the diaphragm in synchrony with mechanical ventilation. Secondary objectives were to confirm that there was synchrony between inspiration and electrical stimulation, to assess the effect of stimulation on Work of Breathing (WOB), and to determine the effect of stimulation on an accelerometer signal.

Stimulation was delivered with a symmetrical biphasic pulse with the charge density of the lead electrodes limited to 25μC/cm^2^ per phase This level of stimulation has been shown to be safe for neurostimulation and is in line with existing commercial phrenic nerve stimulation devices (11). The stimulator was limited to delivering 25mA at 25Hz with a 300 μs pulse width. The charge density of the Ambu needles exceeded the charge density limit of 25μC/cm^2^ per phase since the exposed electrode tip of the Ambu needle had a significantly smaller surface area.

Ventilation parameters measured throughout the procedure included mode of ventilation, inspiratory time, set pressure or flow rate, plateau time, exhaled tidal volume, set and measured breaths per minute (BPM), Positive end expiratory pressure (PEEP), Fraction of inspired oxygen (FiO_2)_ and trigger flow. Additional parameters measured during the procedure included attributes of lung mechanics, specifically, lung compliance (mL/cmH_2_O) and resistance (cmH_2_O/Lps), and blood gasses. Current, frequency and pulse width associated with the electrical stimulation of the left and right electrical nerves were also logged once the initial stimulation settings and associated adjustments had been made.

### Instrumentation Setup

Figure 3 shows the PEPNS System set-up used for the study. LabChart software (ADInstruments, Colorado Spring, CO) and a PowerLab device (ADInstruments, Colorado Spring, CO) were used to perform data acquisition during the study. Accelerometers were placed to record both leg and diaphragm acceleration/movement during electrical stimulation. The PEPNS Console measured WOB and data was output as an analog signal to the PowerLab and LabChart. Wye flow, wye pressure, detection of inspiration and expiration, and the stimulation signals were recorded at 1000 Hz. The wye flow and pressure were measured using a SpiroQuant H flow sensor (EnviteC-Wismar GmbH, Wismar, Germany) using pressure sensors (HSCDRRN002NDAA3 and SSCMRRN160MDAA3, Honeywell, Charlotte, North Carolina). Flow was corrected for barometric pressure and inspired gas composition to body temperature and pressure saturated. A Puritan Bennett™ 840 ventilator (Medtronic, Minneapolis, MN) was used to provide ventilation during the study. The animals’ lung compliance and resistance were measured before stimulation was initiated and these values were entered into the PEPNS user interface along with barometric pressure, ambient temperature, type of humidification (HME) and the percentage of oxygen being delivered by the ventilator to account for the differences these settings cause in gas density.

**Fig 3.**
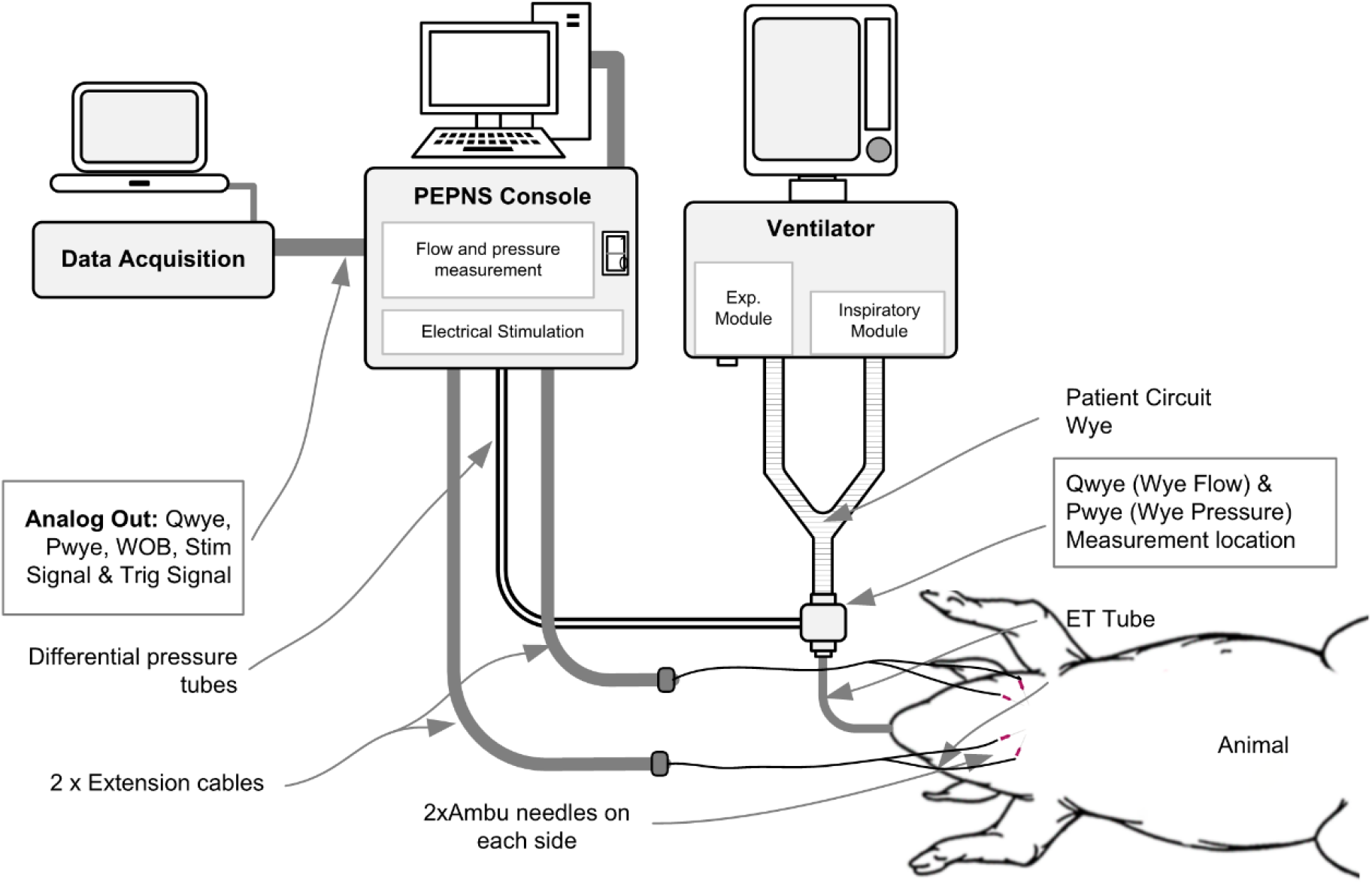
PEPNS System Set-up for Studies.

### Animal Setup, Sedation and Maintenance

The animals were initially sedated with a mixture of Telazol and Xylazine administered intramuscularly at a dose of 3.5 to 5.5 mg/kg followed by Isoflurane gas in 100% O_2_ at 0 to 5% after placement of an endotracheal tube. Isoflurane was only used during the preparation phase of the protocol. Isoflurane was not administered while the ventilator was being used to control breaths, ensuring WOB measurements could be attributed to stimulation only. This ventilation regime required careful monitoring and adjustment to prevent animal inspiratory effort. The animal was brought into a surgical operating room and placed in a supine position on the table. To help maintain a satisfactory body temperature during the treatment procedure, the animals were kept warm with a hot water pad, blankets, and/or a forced-air warming unit. A Foley urinary catheter was placed for drainage of the bladder during the treatment procedure and furosemide was administered at 2 to 4 mg/kg to alleviate any fluid retention issues. Atropine was administered IM at 0.03 to 0.06 mg/kg to decrease respiratory secretions. Isotonic maintenance fluid was administered at a rate of approximately 1 to 2 ml/kg/hour over the course of the 8-hour anesthesia. Intravenous drug infusions administered to the animals during the treatment procedure consisted of ketamine at ~2 mg/kg/hour, Midazolam at ~0.6 mg/kg/hour, and fentanyl ~0.004 mg/kg/hour. Boluses of the drug infusions were given as supplemental anesthetic when immediate attention to maintain surgical plane of anesthesia was needed. Animal vitals such as oxygen saturation and heart rate were recorded every 15 to 60 minutes and arterial or venous blood gas and electrolytes were recorded at baseline and at least once every 3 to 6 hours throughout the treatment procedure.

### Femoral Nerves (Control and Test Group)

For the test and control group, ultrasound imaging was used to successfully identify the position and depth of the left and right femoral nerves prior to lead placement. Femoral nerve depths (S2 Table) were determined during initial needle insertion and at necropsy. The Pajunk needle along with Pajunk stimulation catheter were primed with saline and used to identify the femoral nerve using electrical stimulation with the tip of the needle and catheter used in a bipolar configuration. Once the nerve had been identified by visual confirmation of leg movement, the tip of the needle was inserted past the nerve, facilitating the positioning of the lead electrodes on both sides of the nerve. The Pajunk catheter was then removed while priming the needle with saline as the catheter was retracted. The Pajunk needle was used to place four-pole disposable leads in the area of the left and right femoral nerves. The Pajunk needles were removed by sliding the needle over the lead. An electrical evaluation test was performed to confirm capture was still possible with the deployed lead. Once capture was confirmed, the leads were secured using surgical glue at the lead insertion site and looping the PEPNS lead under a Tegaderm pad to provide tensile relief.

Electrical stimulation optimization was performed on the leads using both leg acceleration signals and visual conformation to determine the pair of electrodes which provided the lowest threshold stimulation currents while maintaining WOB within defined limits. After the completion of the 8-hour electrical stimulation period, the femoral leads were removed using a force gauge to measure the extraction peak and continuous pull force (S1 Table).

### Phrenic Nerve Stimulation (Test Group)

For the test group, a transcutaneous handheld stimulator (SunStim™ Plus, SunMed, Grand Rapids, Michigan) was used to stimulate the left and right phrenic nerves in order to identify their location in the neck region. An ultrasound transducer was then used to identify the location of the carotid artery anatomical landmark and external/internal jugular veins in order to avoid these vessels when inserting and positioning Ambu needles near the left and right phrenic nerves. The Ambu needles were placed using the previously identified landmarks in neck/thoracic inlet region near the left and right phrenic nerve in order to achieve stimulation of the diaphragm.

Once positioned the Ambu needles were secured using surgical glue and diaphragm threshold testing was performed. The minimum electrical stimulation threshold and failure threshold was determined, and stimulation was set at 20% higher than the minimum threshold for the entirety of the 8-hour stimulation period. The animals remained anesthetized and were bilaterally electrically stimulated on each inspiratory breath for up to 8 hours. Accelerometers were placed on the stomach of the animals just below the diaphragm to ascertain the effect of stimulus on the diaphragm. Stimulation frequency during inspiration was set to achieve a desired WOB level between 0.3 and 0.7 joules/L. WOB was measured using the equation of motion, an estimation of the lung compliance and resistance and a measurement of the inspired flow, volume and pressure at the inlet to the endotracheal tube (12).

Test data was collected for each stimulated breath during the first and last hour of stimulation to assess synchrony between inspiration and stimulation and to assess effect on WOB with effectiveness of stimulation based upon the number of breaths that were stimulated. The diaphragm threshold testing was performed again near the end of the 8-hour stimulation period. After the completion of the 8-hour of testing, the leads were removed, and the animal was recovered.

### Follow-up Assessment

Animals were maintained for 29 to 30 days post-treatment and assessed on a daily basis for overall health and moribundity. The animals were then terminated and underwent gross pathological evaluation of the percutaneous insertion sites and diaphragm tissue was collected for histopathological evaluation. Histopathologic evaluation was performed by a pathologist and included qualitative and semi-quantitative morphological observations to assess the biological response of the nerve to the study conditions. These observations included, but not limited to, assessment of the extent and/or severity, with or without grading, of inflammation, necrosis, fibrin, hemorrhage, fibrosis, and nerve fiber degeneration.

### Data Analysis

#### Determination of Sample Size

The sample size for the study was based on the performance endpoint of the ability of the stimulation pulse to capture the diaphragm and evaluation of the following statistical hypothesis:

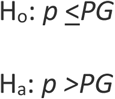

where *p* is the proportion of capture and *PG* is the performance goal of 90%.

The sample size for the performance endpoint was determined based on the assumptions of a power of 80%, one-sided alpha of 0.025, a performance goal of 90%, and an assumed success rate of 97.5%. Based on these assumptions, a minimum of 80 observations were required for at least 80% power to meet the performance goal. For the purposes of sample size calculations, the observations were assumed to be independent; however, the primary analysis methods were planned to account for within animal correlation due to repeated measurements.

#### Statistical Analyses

In general, continuous data were summarized descriptively with mean, standard deviation, and number of evaluable observations with 95% confidence interval based on the t-distribution, where appropriate. Categorical variables were summarized with frequency counts and percentages. All statistical hypothesis tests were performed at the two-sided alpha of 0.05 significance level unless otherwise noted. All analyses were conducted using SAS v9.4 (SAS Institute Inc., Cary, NC, USA).

The performance endpoint was defined as the stimulation pulse to capture the diaphragm and was assessed using accelerometer measurements on the chest to determine if stimulation had a physical effect on tile diaphragm. Successful capture of the diaphragm was determined based on the alignment of the stimulation zone and the accelerometer (aligned=success, not aligned=failure). Each breath was considered captured if the WOB value was between 0.3 and 0. 7 J/L. The rectified and filtered acceleration signal also had to be above a specific threshold level. The performance endpoint was assessed with the following hypothesis:

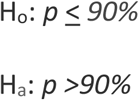

where *p* is the proportion of capture and *PG* is the performance goal of 90%.

The hypothesis was evaluated using the lower bound of a two-sided 95% Wald confidence interval for proportion of successful capture as estimated from a generalized linear random effects model. The endpoint was considered to be met if the lower bound of the two-sided 95% confidence interval was greater than 90%. The generalized linear model was fit with capture as the outcome with random effects for time period (first hour and last hour) and animal to account for multiple measurements within animal and time period. The estimate of the proportion of successful capture and confidence interval were estimated as the Logit-transform of the log odds for capture and corresponding 95% Wald confidence interval.

Inspiration and expiration trigger times were summarized descriptively by animal and time period (first and last hour) and overall to determine synchrony. Post-hoc statistical hypothesis testing was performed for the average inspiration and expiration trigger times at each time period and combined across time periods, separately. The following post-hoc hypothesis was evaluated:

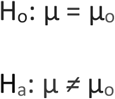

Where μ is the mean trigger in seconds and μ_o_=0.088s for inspiration trigger times and μ_o_ = −0.088s for expiration trigger times.

For each time period and overall, the inspiration and expiration trigger times were presented descriptively with two-sided 95% confidence intervals based on the t-distribution and p-value from a two-sided one-sample t-test. As the hypothesis testing was post-hoc, no adjustments for multiplicity were made. In a post-hoc analysis, capture was summarized based on work of breathing and the accelerometer separately for each animal and overall with simple summary statistics and exact 95% confidence interval.

## RESULTS

All treatment procedures were successful and there were no treatment procedure related complications. Of the six animals treated on this study, four survived until the scheduled termination date. The two animals that did not survive were enrolled in the control arm of the study and their deaths were found not to be related to the study procedures. One animal died due to complications related to a twisted duodenum with adhesions to the stomach and liver. The other animal was terminated early due to due to severe respiratory complications related to a laryngotracheal necrotic lesion. Table 1 lists pre-treatment and pre-termination data for all 6 animals in the study.

The performance endpoint of percentage of successful capture determined by WOB being stimulated within the range of 0.3 to 0.7 joules/L was 99.41%, with the lower bound of the two-sided 95% confidence interval of 95.14%. These results exceeded the prespecified performance goal of 90%. The discordance in capture between WOB and the accelerometer was less than 2%. Table 2 lists the mean inspiration and expiration trigger times for the first and last hours and both hours combined. Table 3 shows the mean WOB for the same time periods. A post-hoc analysis showed mean inspiration trigger times were statistically significantly less than 0.088 seconds for all time points and the mean expiration trigger times were statistically significantly less than −0.088 seconds for all time points. WOB was similar during both the first and last hour observation periods.

**Table 2.** Mean Inspiration and Expiration Trigger Times.

| Time Period | Variable | Animal<br>17P0923 | Animal<br>17P0981 | Animal<br>17P1001 | Mean ± Std<br>(n) | 95%<br>Confidence<br>Interval<sup>1</sup> | P-value<sup>2</sup> |
| --- | --- | --- | --- | --- | --- | --- | --- |
| First Hour | Insp Trigger Time (s) | 0.025 ± 0.016<br>(53) | 0.019 ± 0.015<br>(53) | 0.015 ± 0.004<br>(54) | 0.019 ± 0.013<br>(160) | 0.017, 0.021 | <.0001 |
|  | Exp Trigger Time (s) | -0.032 ± 0.019<br>(53) | -0.008 ± 0.019<br>(53) | 0.003 ± 0.007<br>(54) | -0.012 ± 0.022<br>(160) | -0.015, -0.009 | <.0001 |
| Last Hour | Insp Trigger Time (s) | 0.037 ± 0.015<br>(53) | 0.015 ± 0.003<br>(53) | 0.014 ± 0.004<br>(55) | 0.022 ± 0.014<br>(161) | 0.02, 0.024 | <.0001 |
|  | Exp Trigger Time (s) | -0.021 ± 0.018<br>(53) | 0 ± 0.004 (53) | 0.001 ± 0.004<br>(55) | -0.007 ± 0.015<br>(161) | -0.009, -0.004 | <.0001 |
| Both Hours<br>Combined | Insp Trigger Time (s) | 0.031 ± 0.017<br>(106) | 0.017 ± 0.011<br>(106) | 0.015 ± 0.004<br>(109) | 0.021 ± 0.014<br>(321) | 0.019, 0.022 | <.0001 |
|  | Exp Trigger Time (s) | -0.027 ± 0.019<br>(106) | -0.004 ± 0.014<br>(106) | 0.002 ± 0.006<br>(109) | -0.009 ± 0.019<br>(321) | -0.011, -0.007 | <.0001 |

Data presented as mean ± std (n = sample size)
<sup>1</sup> Confidence interval for mean of all animals combined
<sup>2</sup> P-value based on a two-sided t-test comparing the mean to 0.088 secs for Insp time and -0.088 secs for Exp time.

**Table 3.** Mean Work of Breathing (WOB)

| Time Period | Summary | Animal 17P0923 | Animal 17P0981 | Animal 17P1001 | All Animals |
| --- | --- | --- | --- | --- | --- |
| First Hour | Mean ± Std (n) | 0.396 ± 0.036 (1020) | 0.429 ± 0.054 (1080) | 0.560 ± 0.044 (1202) | 0.467 ± 0.085 (3302) |
|  | Range | (0.286, 0.588) | (0.251, 0.588) | (0.383, 0.691) | (0.251, 0.691) |
|  | 95% Confidence Interval | (0.393, 0.398) | (0.426, 0.433) | (0.558, 0.563) | (0.464, 0.470) |
| Last Hour | Mean ± Std (n) | 0.390 ± 0.032 (1021) | 0.419 ± 0.095 (1080) | 0.513 ± 0.084 (1201) | 0.444 ± 0.093 (3302) |
|  | Range | (0.288, 0.469) | (-0.261, 0.841) | (0.275, 0.956) | (-0.261, 0.956) |
|  | 95% Confidence Interval | (0.388, 0.392) | (0.413, 0.424) | (0.508, 0.518) | (0.441, 0.447) |
| Both Hours Combined | Mean ± Std (n) | 0.393 ± 0.034 (2041) | 0.424 ± 0.077 (2160) | 0.537 ± 0.071 (2403) | 0.455 ± 0.090 (6604) |
|  | Range | (0.286, 0.588) | (-0.261, 0.841) | (0.275, 0.956) | (-0.261, 0.956) |
|  | 95% Confidence Interval | (0.391, 0.394) | (0.421, 0.427) | (0.534, 0.540) | (0.453, 0.458) |

Data presented as mean ± std (n = sample size)
Expected range of WOB 0.3 to 0.7 J/L (joules/liter)

Results of capture from WOB and the accelerometer are summarized in Table 4 and displayed graphically in Figure 4. The accelerometer captured 100% (95% Cl: 99.9%, 100.0%) of all breaths while WOB captured 98.5% (95%CI: 98.2%, 98.8%) of all breaths. The discordance between WOB and Accelerometer was low with only 1.5% (95% Cl: 1.2%, 1.8%).

**Table 4.** Capture by WOB and Accelerometer.

| **Animal ID** | **Method** | **Proportion of Success, % (n/N)** | **95% Confidence Intervals** |
| --- | --- | --- | --- |
| 17P0923 | WOB | 99.8% (2037/2041) | 99.5, 99.9 |
|  | Accelerometer | 100.0% (2041/2041) | 99.8, 100.0 |
| 17P0981 | WOB | 96.8% (2090/2160) | 95.9, 97.5 |
|  | Accelerometer | 100.0% (2160/2160) | 99.8, 100.0 |
| 17P1001 | WOB | 99.1% (2381/2403) | 98.6, 99.4 |
|  | Accelerometer | 100.0% (2403/2403) | 99.8, 100.0 |
| All animals | WOB | 98.5% (6508/6604) | 98.2, 98.8 |
|  | Accelerometer | 100.0% (6604/6604) | 99.9, 100.0 |

**Fig 4.**
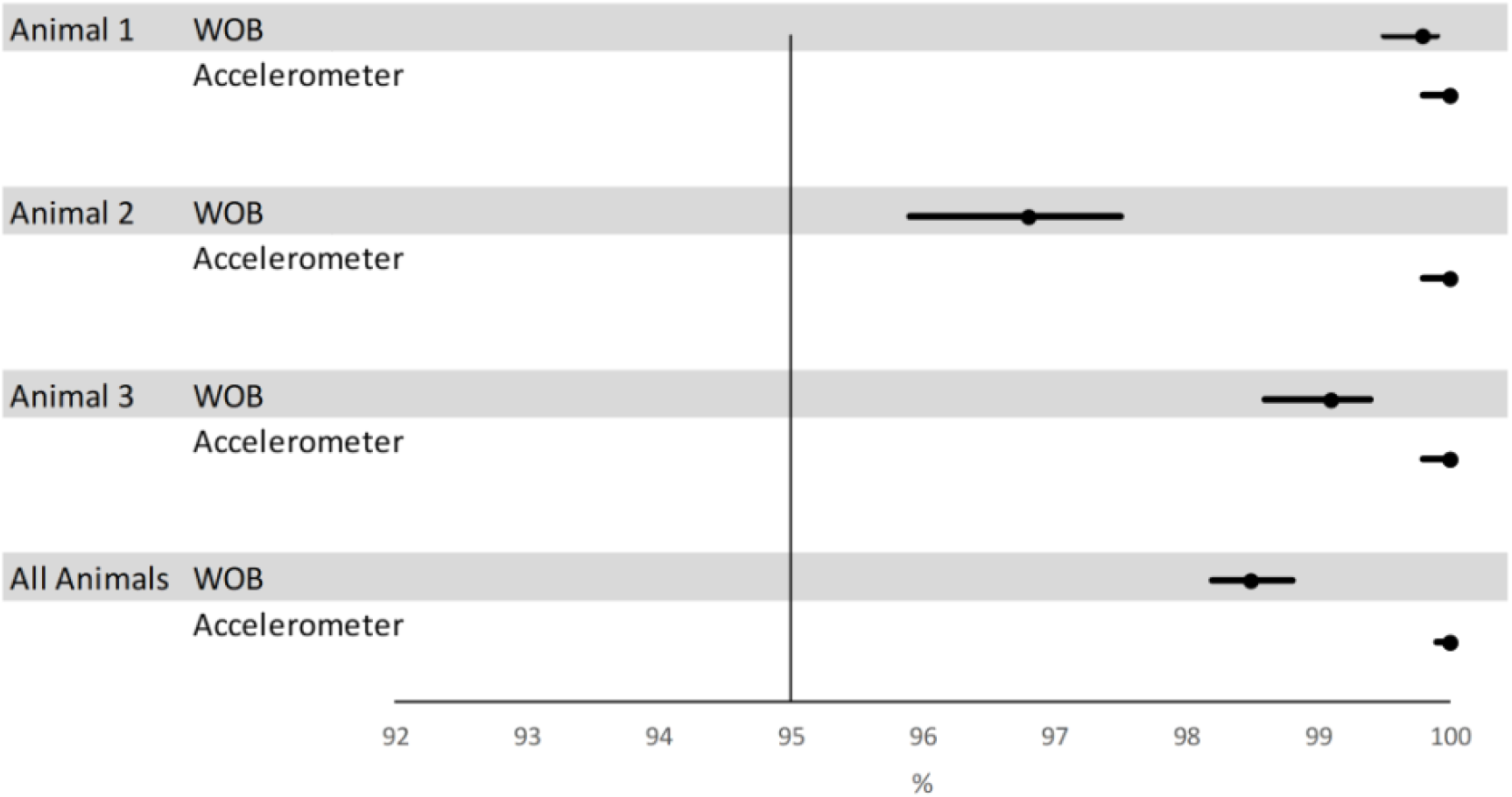
Capture by WOB and Accelerometer.

All surviving animals utilized for study analysis were generally healthy throughout the duration of the study. Gross necropsy of the animals revealed there were no abnormal findings in the majority of organs examined that could have been related to the electrical stimulation treatment procedure. There were no injuries to organs adjacent to the treatment procedure regions, no infections associated with the treatment procedure, no lameness or difficulty breathing, nor any procedure related adverse events. Organs with abnormal findings at necropsy revealed only background findings occasionally seen in swine. Histopathological evaluation also did not reveal any abnormal findings of the right and left femoral nerves or the diaphragm that could have been related to the electrical stimulation treatment procedure.

## DISCUSSION

The two evaluation methods described in the present study provided important pre-clinical information on the safety and efficacy of percutaneously placed proprietary multipole lead and the PEPNS system prior to planned human clinical studies. We confirmed the ability to safely place the leads using ultrasound imaging and to maintain capture over 8 hours without causing femoral nerve injury. Importantly, the study demonstrated that electrical stimulation of the phrenic nerve did not produce diaphragm injury as noted by the negative histopathological findings and the lack of treatment associated injuries. It also achieved the capture performance endpoint by demonstrating stimulation was synchronous with inspiration over the 8 hours of continuous stimulation. The above findings collectively indicated that the electrode placement procedure and the associated stimulation protocol would be transferable to humans in a clinical setting. The results of this GLP study served as the basis for developing the clinical protocol for the recently completed PEPNS first-in-human clinical study [6].

Multiple animal models (caprine, ovine, canine, and porcine) were assessed prior to the present study to determine the most appropriate animal model which could simulate the neck anatomy of humans prior to our decision to use a porcine model. Several authors have reported on the use of porcine models for either acute and chronic phrenic nerve pacing and to study the effect of prolonged mechanical ventilation on diaphragmatic function [13–15]. While there are no quadruped species that have a complete phrenic nerve outside of the thoracic cavity, the porcine model used for the present study was deemed suitable since ruminants in general are not good subjects for general anesthesia due to their anatomy and physiological responses being less predictable [16]. Additionally, the danger of regurgitation and inhalation of ingesta is much greater in ruminant species compared to other species. Swine are known to have similar dermatologic and neurologic responses to that of the human when utilized for medical device testing [17]. As a result, a swine model was determined to be most appropriate for the safety evaluations performed as a part of the present study.

The femoral nerve was successfully captured during this study using 4-electrode lead at the start and end of an 8-hour test period without the need to increase the amplitude of the current delivered to the nerve or reposition the lead. This 0.87 mm diameter lead was designed to enable minimally invasive percutaneous insertion at the patient bedside, avoiding the need for more invasive procedural approaches for lead placement in order to stimulate the phrenic nerve. The demonstrated ability to easily place the leads without local injury in the porcine model utilized for this study supports the rationale for using these leads in humans and the potential for ease of insertion and safety when placed in a patient’s neck region.

The decision to deliver electrical stimulation to the phrenic nerve over an 8-hour period to contract the diaphragm was chosen to increase the likelihood of animal survivability and was deemed to be more than sufficient to demonstrate the safety of the initial proposed intermittent stimulation for a shorter duration over a 48-hour period for planned first in human studies. Previous preclinical testing had demonstrated that it would be difficult to survive animals after prolonged periods of general anesthesia with the determinant of survivability being duration of sedation as opposed to the duration of electrical stimulation.

A trigger time of 88ms for as a threshold for determining synchrony between stimulation and the beginning of both inspiration and expiration would be consider fast for inspiratory breath detection. Chen et al. reported that inspiratory triggering is greater than 125ms for all ventilators they examined with expiratory triggering delay times having similar or larger ranges (18). Thille et al. also reported that significant differences were found across ventilators with a range of inspiratory triggering delay from 42ms to 88ms (19).

Limitations of the present study include the lack of randomization and the use of an unblinded study design which could potentially introduce bias into the interpretation of the results. Also, while porcine anatomy and physiology are similar to those of humans, the lack of collateral ventilation in pigs and the impact of this on respiratory function [20) suggests drawing clinical conclusions from this study relative to lung function should be avoided.

## CONCLUSION

With respect to the assessment of the safety of percutaneous electrical stimulation of the phrenic to contract the diaphragm during inspiration and ultrasound guided placement of the a multipole lead beside the femoral nerves, the findings related to tissue response, animal health, and stimulation effect endpoints of this study indicate the safety of the device used in this porcine model.

## Acknowledgements

The authors would like to thank James Berry from Gallivant Medical Imaging, Hudson, Wisconsin, for his assistance with performing the ultrasound imaging and needle insertions for this study. Medical writing assistance was provided by Larry Yost from The Atticus Group, LLC (Portsmouth, New Hampshire, USA).

## DECLARATIONS

### Funding

This study was funded by a research grant from Stimdia Medical, Inc.

### Ethics approval

All studies were carried out in an Association for Assessment and Accreditation of Laboratory Animal Care (AAALAC) approved facility (American Preclinical Services, LLC, Minneapolis, Minnesota) and were reviewed and approved by the Institutional Animal Care and Use Committee (IACUC).

### Authors’ contributions

JO, CD, and JM designed the study. JO, CD and IP performed the experiments. JM analyzed the data. JO and TJ wrote the manuscript. TJ critically reviewed and updated the manuscript based on author feedback. All authors approved the final version of the manuscript.

## Notes

### Competing Interest Statement

JOM and TJ are employees of Stimdia Medical. CD and IP are employees of American Preclinical Services, LLC who was contracted to perform this study by Stimdia Medical. JM is an employee of NAMSA who was contracted by Stimdia Medical to perform the statistical analysis for the present study.

